# Fusion of Video and Inertial Sensing Data via Dynamic Optimization of a Biomechanical Model

**DOI:** 10.1101/2022.11.15.516673

**Authors:** Owen Pearl, Soyong Shin, Ashwin Godura, Sarah Bergbreiter, Eni Halilaj

## Abstract

Inertial sensing and computer vision are promising alternatives to traditional optical motion tracking, but until now these data sources have been explored either in isolation or fused via unconstrained optimization, which may not take full advantage of their complementary strengths. By adding physiological plausibility and dynamical robustness to a proposed solution, biomechanical modeling may enable better fusion than unconstrained optimization. To test this hypothesis, we fused RGB video and inertial sensing data via dynamic optimization with a nine degree-of-freedom model and investigated when this approach outperforms video-only, inertial-sensing-only, and unconstrained-fusion methods. We used both experimental and synthetic data that mimicked different ranges of RGB video and inertial measurement unit (IMU) data noise. Fusion with a dynamically constrained model significantly improved estimation of lower-extremity kinematics over the video-only approach and estimation of joint centers over the IMU-only approach. It consistently outperformed single-modality approaches across different noise profiles. When the quality of video data was high and that of inertial data was low, dynamically constrained fusion improved estimation of joint kinematics and joint centers over unconstrained fusion, while unconstrained fusion was advantageous in the opposite scenario. These findings indicate that complementary modalities and techniques can improve motion tracking by clinically meaningful margins and that data quality and computational complexity must be considered when selecting the most appropriate method for a particular application.

## 1. INTRODUCTION

Accessible motion tracking could transform rehabilitation research and therapy. The traditional marker-based approach is limited to specialized laboratories equipped with expensive infrared camera optical motion tracking systems and trained personnel. Inertial sensing and standard red-green-blue (RGB) camera computer vision-based approaches offer greater flexibility, given their low cost and portability, but collective understanding of the strengths and weaknesses of kinematics estimation algorithms associated with each technology is still evolving (Table 1). Additionally, efforts to merge the strengths of these complementary technologies are sparse.

**Table 1.** Qualitative comparison of state-of-the-art IMU and video-based motion capture techniques for measuring joint kinematics.

| Modality | Method | Example Articles | Advantages | Disadvantages |
| --- | --- | --- | --- | --- |
| IMUs | Sensor-fusion Filters (e.g., EKF, Madgwick, Mahoney) | Mahony, 2008. Madgwick, 2010. Sabatini, 2011. Joukov, 2014. Al Borno, 2022. | Computationally efficient;<br>Open source | Magnetometers are often unreliable<br>Magnetometer-free approaches are inaccurate |
|  | Deep Learning (e.g., CNNs, LSTMs, Transformers) | Huang, 2018. Rapp, 2021. Yi, 2021-22. Mundt, 2020-2021. | Implicitly learns noise;<br>Open source | Training data are not sufficiently representative of pathologies and activities |
|  | Biomechanical Modeling: Static Optimization | Roetenberg, 2013. Karatsidis, 2016-19. | Predicts GRFs, muscle forces, and joint reaction forces | Requires drift correction using additional sensors;<br>Computational cost;<br>Closed source |
|  | Biomechanical Modeling: Direct Collocation | Dorschky, 2019. | Predicts GRFs, muscle forces, and joint reaction forces | Requires drift correction using limiting assumptions;<br>Computational cost;<br>Closed source |
| Videos | Deep learning & Unconstrained Optimization | Kanazawa, 2018. Isakov, 2019. Zhang, 2020. Kocabas, 2020-21. | Computationally efficient;<br>Open source | Data-driven: training data not representative of clinical populations;<br>Sensitive to occlusions |
|  | Deep Learning & Biomechanical Modeling | Kanko, 2021. Strutzenberger, 2021. Uhlrich, 2022. | Predicts GRFs, muscle forces, and joint reaction forces;<br>Open source | Data-driven: training data not representative of clinical populations;<br>Computational cost |
| IMUs & Videos | Deep learning & Unconstrained Optimization | Halilaj, 2021. | Computationally efficient;<br>Merges complementary modalities;<br>No integration of inertial data necessary | Poor initial estimations from video are propagated in the optimization |
|  | Deep learning & Dynamically Constrained Optimization | Proposed Method | Predicts GRFs, muscle forces, and joint reaction forces;<br>Merges complementary modalities while satisfying the laws of physics<br>No integration of inertial data necessary;<br>Accurate with noisy IMU data | Currently, 2-D proof of concept with 3-D validity remaining to be tested;<br>Computational cost |

Vision-based methods using RGB cameras are successful in camera-dense environments, but occlusion continues to pose challenges in reduced-camera settings (Joo et al., 2019). Although now widely used in robotics applications, translation of vision-based methods to human movement sciences remains uncommon due to accuracy limitations (Seethapathi et al., 2019). Computer vison models are data-driven and typically not constrained to satisfy physiological constraints. Biomechanical modeling has been considered as a possible approach for improving the accuracy of computer vision approaches and making them more accessible to the biomechanics community (Kanko et al., 2021; Strutzenberger et al., 2021; Uhlrich et al., 2022). Although comparisons with marker-based data suggest that the accuracy of these methods ranges widely between 3° – 20°, depending on the degree-of-freedom, no study to date has systematically discerned how this accuracy compares to alternative approaches and to what degree the incorporation of biomechanical models improves results.

Similarly, converting multimodal time series data from inertial measurement units (IMU) into accurate joint kinematics remains challenging due to the many possible sources of uncertainty, including bias noise, thermo-mechanical white noise, flicker noise, temperature effects, calibration errors, and soft-tissue artifacts (Park & Gao, 2008; Picerno, 2017). Traditional sensor fusion filters used to mitigate drift (Madgwick, 2010; Mahony et al., 2008; Sabatini, 2011) typically rely on magnetometers, which are susceptible to ferromagnetic interferences (de Vries et al., 2009). The results of sensor-fusion filters have been refined with biomechanical models (Al Borno et al., 2022), but whether findings will translate to natural environments remains uncertain because marker-based motion capture was used for sensor-to-body calibration, IMUs impacted by ferromagnetic disturbances were manually excluded, and the effect of soft-tissue motion was partly eliminated by attaching IMUs to solid marker cluster plates, helping the IMUs move rigidly with the marker clusters. Deep learning has been proposed as an alternative (Mundt et al., 2020; Rapp et al., 2021) but has been limited by datasets that are not representative of all activities and clinical populations. Constrained optimization via biomechanical modeling, both static and dynamic, has also been used for estimation of both kinematics and kinetics. Static optimization approaches rely on zero-velocity detection algorithms from joint constraints, external contacts, and additional sensors (GPS, RF-based local positioning sensors, barometers, etc.) to correct the position of the model at each step (Karatsidis et al., 2019; Roetenberg et al., 2013), while dynamic optimization approaches currently require that the motion be periodic (Dorschky et al., 2019), both of which limit ease of implementation and generalizability.

IMU and vision data have complementary strengths that can be leveraged to overcome their individual limitations, but it is unclear if fusion via a dynamically constrained biomechanical model would improve estimation of kinematics over unconstrained optimization (Halilaj et al., 2021). Inertial sensing can compensate for occlusions in videos, videos can compensate for drift in inertial data, and biomechanical models can add physiological plausibility and dynamical robustness. Here we fuse video and IMU data via dynamic optimization of a nine degree-of-freedom (DOF) model (Fig. 1) and investigate the circumstances under which this approach outperforms (1) standard computer vision techniques using video data, (2) dynamic optimization of a biomechanical model using IMU data, and (3) fusion of IMU and video data via unconstrained optimization (i.e., without a biomechanical model). In addition to comparing these methods using experimental data, we quantified their sensitivity to IMU and video data noise by scaling each subject’s unique noise backgrounds. We hypothesized that fusion of video and IMU data with biomechanically constrained optimization would improve estimation of kinematics over the alternatives under all the noise profiles. We have shared a MATLAB library to encourage testing of these techniques with additional data and the exploration of new scientific questions.

**Fig. 1.**
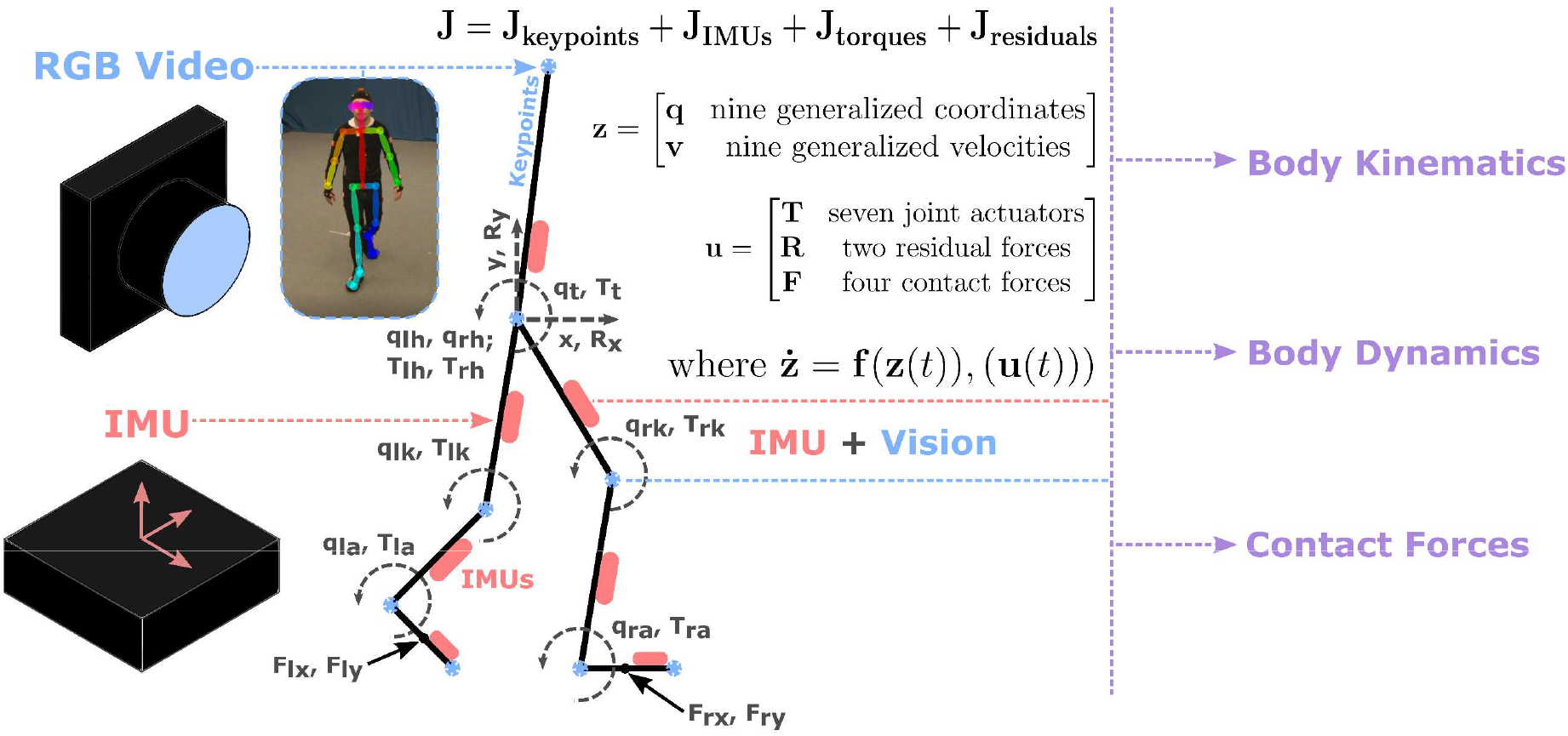
Biomechanical Model and Dynamics Overview. Red-green-blue (RGB) video and inertial measurement unit (IMU) data are fused into a single optimal control trajectory tracking problem, where the state of a planar musculoskeletal model is optimized to produce joint center trajectories and inertial profiles that match the experimental data. A nine degree-of-freedom (two translational, seven rotational) model is actuated by seven joint torques, four ground contact forces, and two residual forces accounting for dynamic inconsistencies due to modeling simplifications. The model fuses data from eight anatomical keypoints acquired from three-dimensional triangulation of computer vision keypoints extracted from RGB video data and seven inertial measurement units placed on each rigid body segment. Direct collocation is used to minimize a cost functional with keypoint and IMU tracking error costs and an effort cost for regulating the joint torques and residual forces.

## 2. METHODS

### 2.1 Biomechanical Model

To test our leading hypothesis, we used a planar nine DoF biomechanical model that consisted of seven rigid body segments (Fig. 1). One segment represented the head, arms, and torso and three segments represented each leg. Body-segment lengths, masses, and mass moment of inertias were estimated by scaling a three-dimensional musculoskeletal model based on 21 cadavers and 24 young adults (Delp et al., 1990, 2007) with marker-based motion capture data. The model state, ***z***, contained nine generalized coordinates, ***q***, and their generalized velocities, ***v***, consisting of the horizontal and vertical sagittal plane translation of the pelvis, *x* and *y*, and the sagittal plane rotation (flexion-extension angles) of the pelvis, hip joints, knee joints, and ankle joints, *q_t_*, *q*_lh_, *q*_rh_, *q*_lk_, *q*_rk_, *q*_la_, *q*_ra_, respectively:

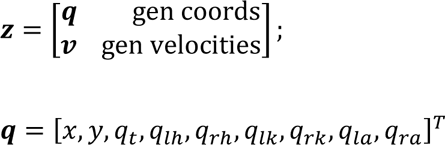

The model control vector, ***u***, contained joint torques, ***T***, contact forces, ***F***, and residual forces accounting for dynamic inconsistencies due to modeling simplifications, ***R*** (Fig. 2c). Residual forces are artificial forces applied at the pelvis of the model to help position and orient it in space when real forces alone are insufficient due to modeling simplifications.

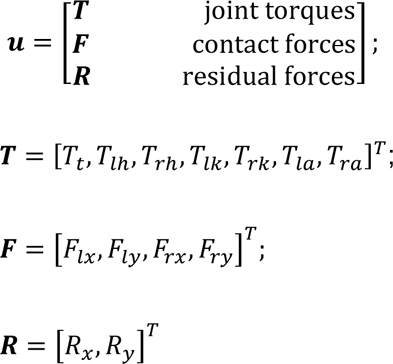

**Fig. 2.**
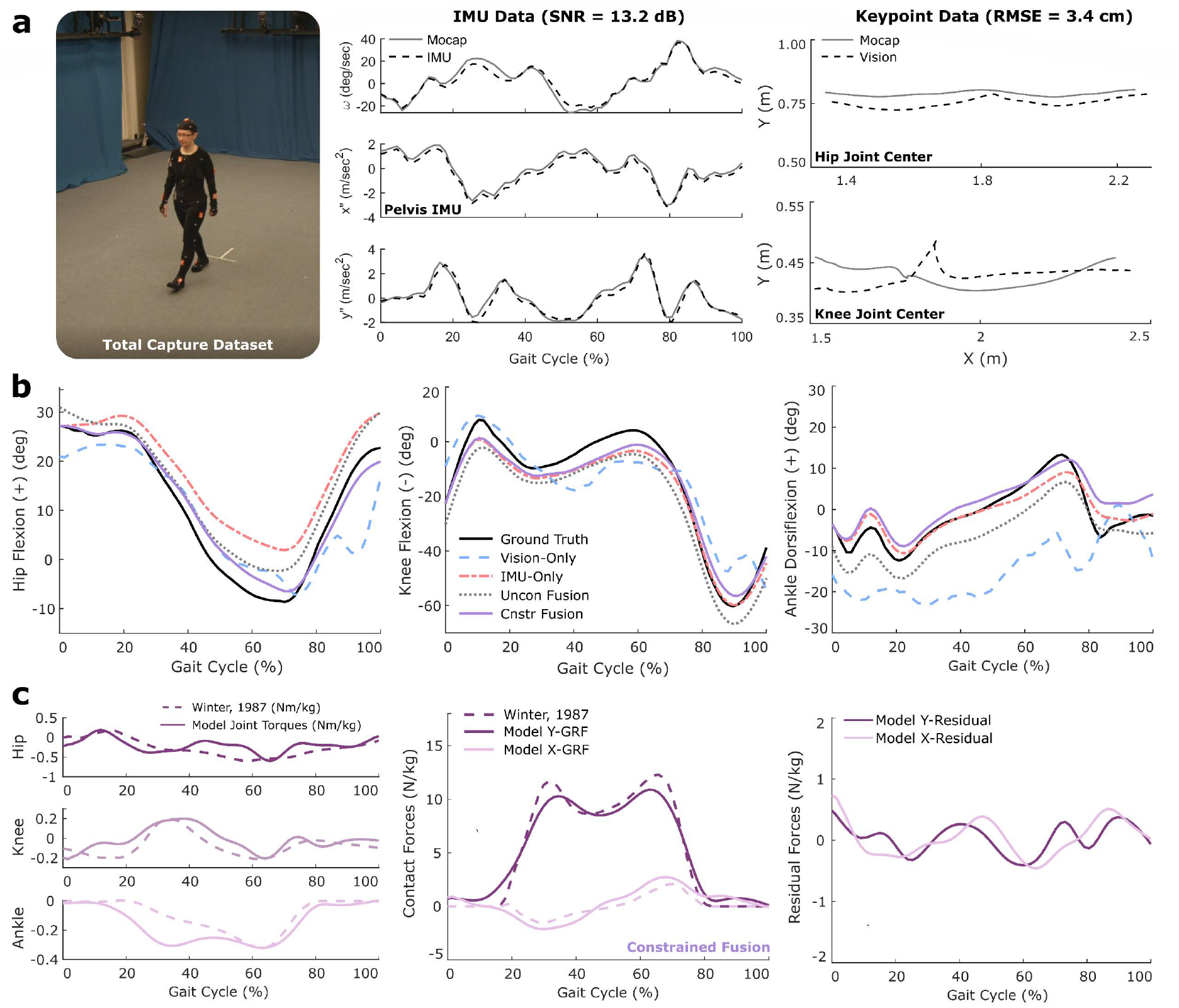
Experimental Data and Resulting Markerless Kinematics. **(a)** We used walking data from 5 subjects recorded with marker-based motion capture, inertial measurement units (IMU), and four videos. The IMU data had a signal-to-noise (SNR) ratio of 13.2 dB, while the video-based keypoints (i.e., joint center positions) had a root-mean-squared error (RMSE) of 3.4 cm. **(b)** Dynamically constrained fusion of IMU and video data via a biomechanical model and direct collocation (Cnstr Fusion, in solid magenta) improved kinematic predictions over competing markerless motion capture approaches (shown for a single female subject). **(c)** Although estimation of kinetics was not a study goal, constrained fusion estimated contact forces (middle) and joint torques (left) relatively well (Winter, 1987). Residual forces (right) are non-physical forces that account for discrepancies between the model and reality when modeling assumptions cause inconsistencies between the biomechanical model and the subject’s true multibody dynamics.

We used Autolev (Symbolic Dynamics Inc; Sunnyvale, CA) and Kane’s equations of motion to derive symbolic expressions for the nine equations of motion in their explicit form and implemented them in MATLAB (Mathworks, Inc; Natick, MA):

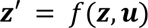

### 2.2 Experimental Data

To test the four markerless approaches for predicting joint kinematics, we used overground walking data from five subjects (4 male; 1 female) from Total Capture (Fig. 2a), a publicly available dataset commonly used to benchmark computer vison methods for motion tracking (Trumble et al., 2017). Motion was captured in a 4 x 6 m area with eight high definition (HD) RGB video cameras at 60 Hz, seven Xsens IMUs (Xsens; Enschede, The Netherlands) positioned on the pelvis, left and right thigh, left and right shank, left and right foot at 1000 Hz, and an infrared camera marker-based motion capture system (Vicon Industries, Inc; Hauppauge, NY) at 100 Hz. Sagittal plane projections of the video and IMU data were used as inputs for the biomechanical model.

### 2.3 Kinematics Estimation: Vision-Only

We extracted two-dimensional (2-D) keypoints (i.e., joint centers) and the confidence score associated with each keypoint from each RGB video camera using the Cascaded Pyramid Network (CPN) (Chen et al., 2018). We triangulated the keypoints by using a direct linear transformation algorithm to extract three-dimensional (3-D) keypoints (Hartley & Sturm, 1997). Contributions from each video were weighted by the confidence score associated with the corresponding 2-D keypoint. We computed kinematics by minimizing the error between the triangulated keypoints derived from video data and the joint centers of the biomechanical model.

### 2.4 Kinematics Estimation: Dynamically Constrained Fusion

Our proposed approach fuses RGB video and IMU data by finding the model states ***z***(*t*) and controls ***u***(*t*) over time, such that the simulated keypoint locations and body segment accelerations and angular velocities from the model state match those obtained from experimental video and IMU data. This was done by formulating the following optimal control problem and solving it via direct collocation:

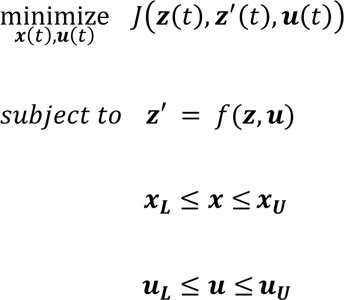

The cost functional *J*(***z***(*t*), ***z***^’^(*t*), ***u***(*t*)) is minimized with respect to a bounded state and control and a constraint on the first derivative of the state vector from the explicit form of the equations of motion. The cost functional includes a tracking term for both the keypoints and the inertial data, *J*_track_, as well as an effort term for both the joint torque actuators and the residual forces, *J*_effort_:

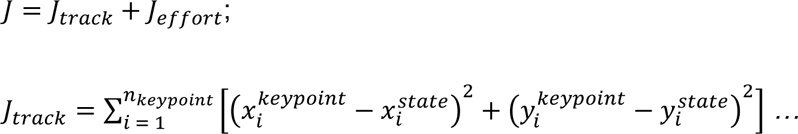

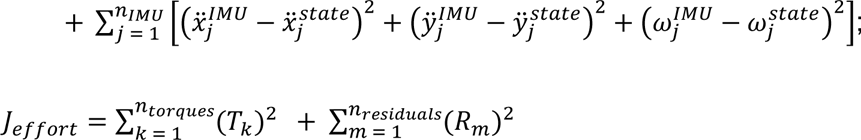

We transcribed the large-scale, sparse nonlinear optimization problem via direct collocation using the OptimTraj library for MATLAB (Kelly, 2017).

### 2.5 Kinematics Estimation: IMU-Only

To perform dynamic optimization with IMU data alone, we took the same steps as in the dynamically constrained fusion approach (2.4) but removed the keypoint terms from within the *J*_track_ portion of the cost. We followed a previously proposed method and applied the assumption that motion was periodic to overcome the drift resulting from integrating noisy IMU data (Dorschky et al., 2019). This involved segmenting the walking data into individual gait cycles and using the mean gait cycle as the input to the *J*_track_ term.

### 2.6 Kinematics Estimation: Unconstrained Fusion

For fusion of IMU and video data via unconstrained optimization, we formulated a simplified optimization problem where *J*_track_ from the IMU and vision optimization was minimized, excluding *J*_effort_ and constraints on system dynamics and model controls (Halilaj et al., 2021). Here, the optimal set of kinematics was determined by minimizing the error between the experimental IMU and video data and the synthetic IMU and keypoint profiles projected from the subject’s current state.

### 2.7 Synthetic Data Generation

In addition to building simulations with the experimentally captured data, we generated synthetic data to investigate how each of the four approaches responded to changes in noise magnitude. We first estimated the naturally occurring noise background, ***φ***, from the experimental data. Ground truth trajectories for each joint center’s position and each body segment’s accelerations and angular velocities were calculated via marker-based motion capture data and analytic equations formed in Autolev, as noted above. Noise was defined as the difference between the ground-truth trajectories and IMU-based (angular velocity and linear acceleration) or video-based (joint center position) trajectories. We then multiplied this experimental noise background by a scale factor, *S*, to achieve synthetic data with new noise magnitudes, without editing the shape of the experimentally observed noise distribution:

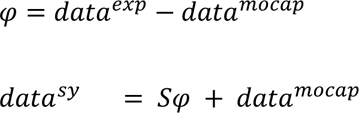

Using infrared camera marker-based motion capture as the ground truth, the mean ± standard deviation keypoint root-mean-square error (RMSE) for the five subjects was 3.5 ± 0.2 cm. We scaled the naturally occurring noise background, ***φ***, for each subject to RMSEs of 6.0, 3.5, and 1.0 cm by adjusting only the scale of the noise background while maintaining its original distribution. These new noise background magnitudes represented low, medium, and high accuracy conditions, based on single-view and multi-view RGB camera approaches (Iskakov et al., 2019; Kadkhodamohammadi & Padoy, 2019; Kanazawa et al., 2018; Kocabas et al., 2020). An RMSE of 6.0 cm corresponds to single-view approaches such as the Human Mesh Recovery (HMR) (Kanazawa et al., 2018) and Video Inference for Body Pose And Shape Estimation (VIBE) (Kocabas et al., 2020). An RMSE of 3.5 cm corresponds to multi-camera algebraic triangulation approaches like what was used in this study. An RMSE of 1.0 cm corresponds to multi-camera methods incorporating learnable triangulation (Iskakov et al., 2019; Kadkhodamohammadi & Padoy, 2019). The IMU data had a mean ± standard deviation signal-to-noise ratio (SNR) of 13.2 ± 0.4 dB.

To generate the IMU synthetic data, we scaled the naturally occurring noise background for each subject to SNRs of 10, 17.5, and 25 dB, which represented low, medium, and high IMU accuracy conditions. These conditions corresponded to IMU data influenced by electrical noise in the form of white noise, scale factor noise, and bias nose (Park & Gao, 2008), a range of commonly occurring static misplacement and misorientation errors (Tan et al., 2019), and a range of previously established soft-tissue motion magnitudes naturally occurring during walking (Fiorentino et al., 2017). To determine appropriate magnitudes to which the experimental IMU noise backgrounds would be scaled, we simulated combinations of misplacement, misorientation, and soft-tissue motion artifacts by formulating analytic equations for each body segment’s accelerations and angular velocity in Autolev:

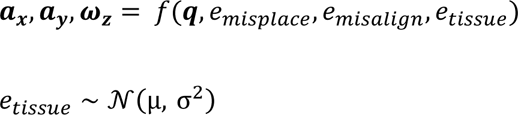

We added error terms while deriving the body segment inertial profiles to model the static misplacement, *e*_misplace_, the static misalignment, *e*_misalign_, and the variable misplacement due to soft-tissue motion, *e*_tissue_. We calculated the noise background magnitudes corresponding to these errors as the difference between the inertial profiles of the body segments derived with and without incorporating the sources of error, and then scaled the error terms to represent the range of expected naturally occurring noise magnitudes (Table 2). We sampled *e*_tissue_ from a normal distribution with μ and σ equivalent to the mean and standard deviation of soft-tissue motion magnitudes measured through X-rays (Fiorentino et al., 2017).

**Table 2.**
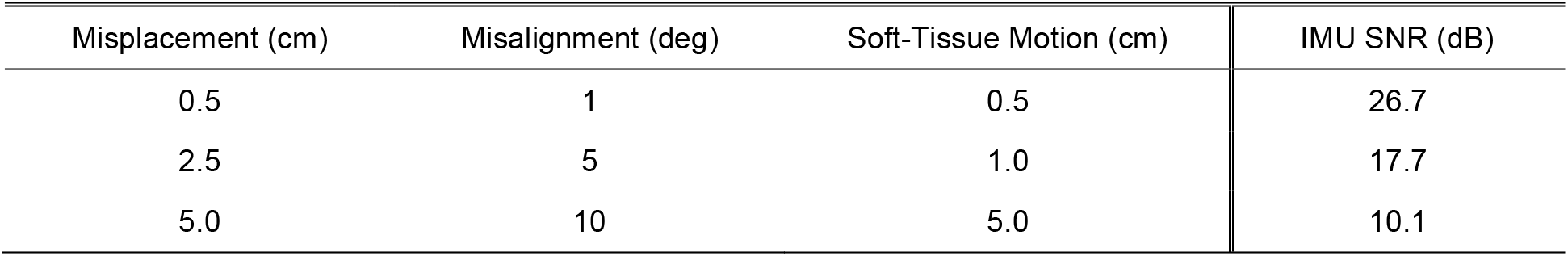
Sources of uncertainty for each modeled IMU signal-to-noise ratio (SNR) profile.

| Misplacement (cm) | Misalignment (deg) | Soft-Tissue Motion (cm) | IMU SNR (dB) |
| --- | --- | --- | --- |
| 0.5 | 1 | 0.5 | 26.7 |
| 2.5 | 5 | 1.0 | 17.7 |
| 5.0 | 10 | 5.0 | 10.1 |

### 2.8 Performance Evaluation

We computed mean-per-joint position error and joint angle error between the simulation results and ground truth infrared camera marker-based motion capture data for each optimization approach and noise profile. We used a one-way repeated measures analysis of variance (RM-ANOVA) and Tukey’s Honest Significant Difference (HSD) for post-hoc analysis to test the leading hypothesis that dynamically constrained fusion would result in lower kinematic errors compared to the other three approaches. The test was carried out for two primary kinematic outcomes: the mean full-body RMSEs for joint angles and joint center positions. A two-way RM-ANOVA followed by an HSD test within noise conditions were used to test the second hypothesis that dynamically constrained fusion would outperform the other three methods when the data were characterized by different noise profiles. The two-way RM-ANOVA considered both the four competing methods and the nine repeated combinations of IMU and video data noise profiles. Results are presented as mean ± standard deviation of the per-joint RMSE compared to marker-based motion capture. An Anderson-Darling test for normality was used to confirm that the data were normally distributed (Yap & Sim, 2011).

## 3. RESULTS

### 3.1 Comparison of Modeling Approaches

Dynamically constrained fusion performed better than single-modality methods, but similarly to unconstrained fusion when using the experimental data (Fig. 2b; Fig. 3). It improved estimation of joint angles by 6.0° ± 1.2° (p < 0.0001) over the vision-only approach and joint centers by 4.5 ± 2.8 cm (p = 0.0018) over the IMU-only approach. Joint angle estimates with the vision-only approach were the least accurate of the four approaches, with RMSEs of 5.1° ± 1.7° for hip flexion, 9.7° ± 3.2° for knee flexion, and 16.0° ± 1.2° for ankle dorsiflexion. Similarly, joint center position estimates with the IMU-only approach were the least accurate of the four approaches, producing errors ranging from 5.6 ± 2.4 cm at the hip to 6.8 ± 3.3 cm at the ankle. The two fusion approaches performed similarly to each other and better than single modality approaches by maintaining accuracy with respect to both joint angles and joint positions. However, dynamically constrained fusion did facilitate improvements over unconstrained fusion in estimates of ankle flexion by 3.3° ± 1.3° (p = 0.0076).

**Fig. 3.**
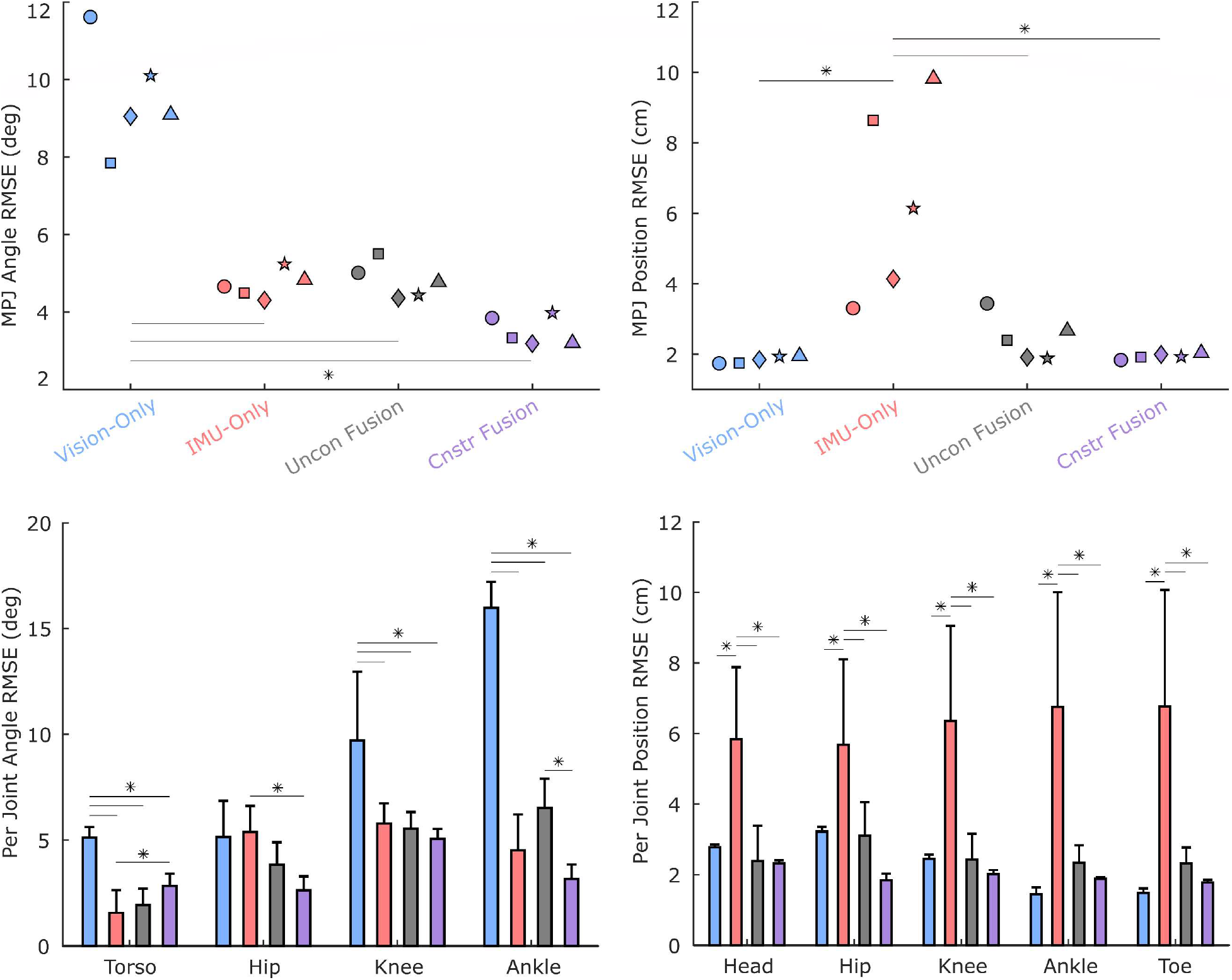
Comparison of Markerless Approaches. Fusion approaches result in lower mean per joint (MPJ) angle root-mean-square errors (RMSEs) (top left) than the vision-only approach and lower MPJ position RMSEs (top right) than the IMU-only approach when tested on experimental data from the Total Capture dataset. Each symbol (top plots) represents a unique subject. Fusion methods resulted in better accuracy than single modality methods by maintaining consistent accuracy with respect to both joint angles and joint center positions across all individual joints. (*p < 0.05)

### 3.2 Sensitivity to Noise

Dynamically constrained fusion performed better than unconstrained fusion when the accuracy of IMU data was low and the accuracy of the video data was high, whereas unconstrained fusion performed better in the opposite scenario (Fig. 4). When the IMU data were of low quality (SNR of 10 dB) and the predicted keypoints from video data were of high quality (RMSE of 1.0 cm), constrained fusion improved estimates of joint angles by RMSEs of 3.7° ± 1.2° (p < 0.0001) and joint centers by 1.9 ± 0.5 cm (p < 0.0001) over unconstrained fusion. When the IMU data were of high quality (SNR of 25 dB) and the predicted keypoints were of low quality (RMSE of 6.0 cm), unconstrained fusion improved estimates of joint angles by 3.0° ± 1.4° (p = 0.0049) and joint center positions by 1.2 ± 0.7 cm (p = 0.0183) over constrained fusion. However, when the quality of IMU data and predicted keypoints was scaled up and down simultaneously, differences between the fusion techniques were not significant.

**Fig. 4.**
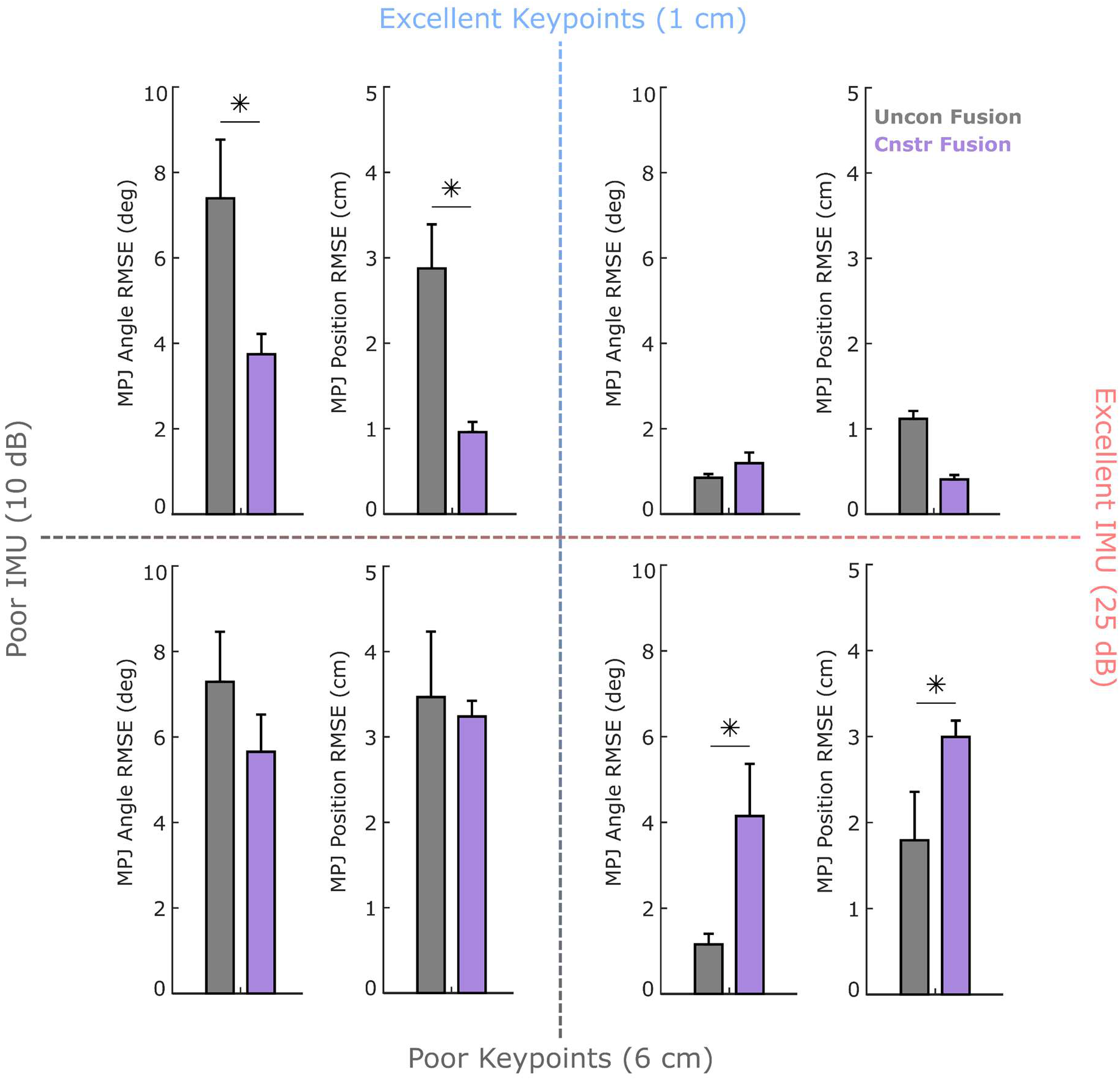
Sensitivity of Fusion Approaches to Noise. Dynamically constrained fusion was advantageous at lower IMU accuracies and higher keypoint accuracies, whereas unconstrained fusion was advantageous at higher IMU accuracies and lower keypoints accuracies. This phenomenon occurs due to the sometimes complementary, but sometimes redundant nature of IMU data and modeling constraints since both provide information on the first and second order derivatives of the body segment motions. Mean ± standard deviation is plotted here with *p < 0.05.

Single-modality approaches generally performed worse than fusion approaches across the varied data qualities, with some exceptions (Fig. 5). The vision-only approach resulted in significantly worse joint angle estimates than the fusion approaches at every condition except when very low IMU data quality (SNR of 10 dB) was paired with very high keypoint data quality (RMSE of 1 cm). At this condition, vision-only matched constrained fusion (p = 0.8071) with a joint angle RMSE of 3.3° ± 0.5°. The IMU-only approach resulted in significantly worse joint center position estimates compared to the fusion approaches at five out of the nine conditions, mainly due to position drift that accumulates over the duration of the IMU-only simulations (Fig. 6). However, at combinations of medium to excellent IMU data accuracy (17.5 – 25 dB) and poor to medium keypoint data accuracy (6.0 – 3.5 cm), the IMU-only approach performed equivalently to fusion methods.

**Fig. 5.**
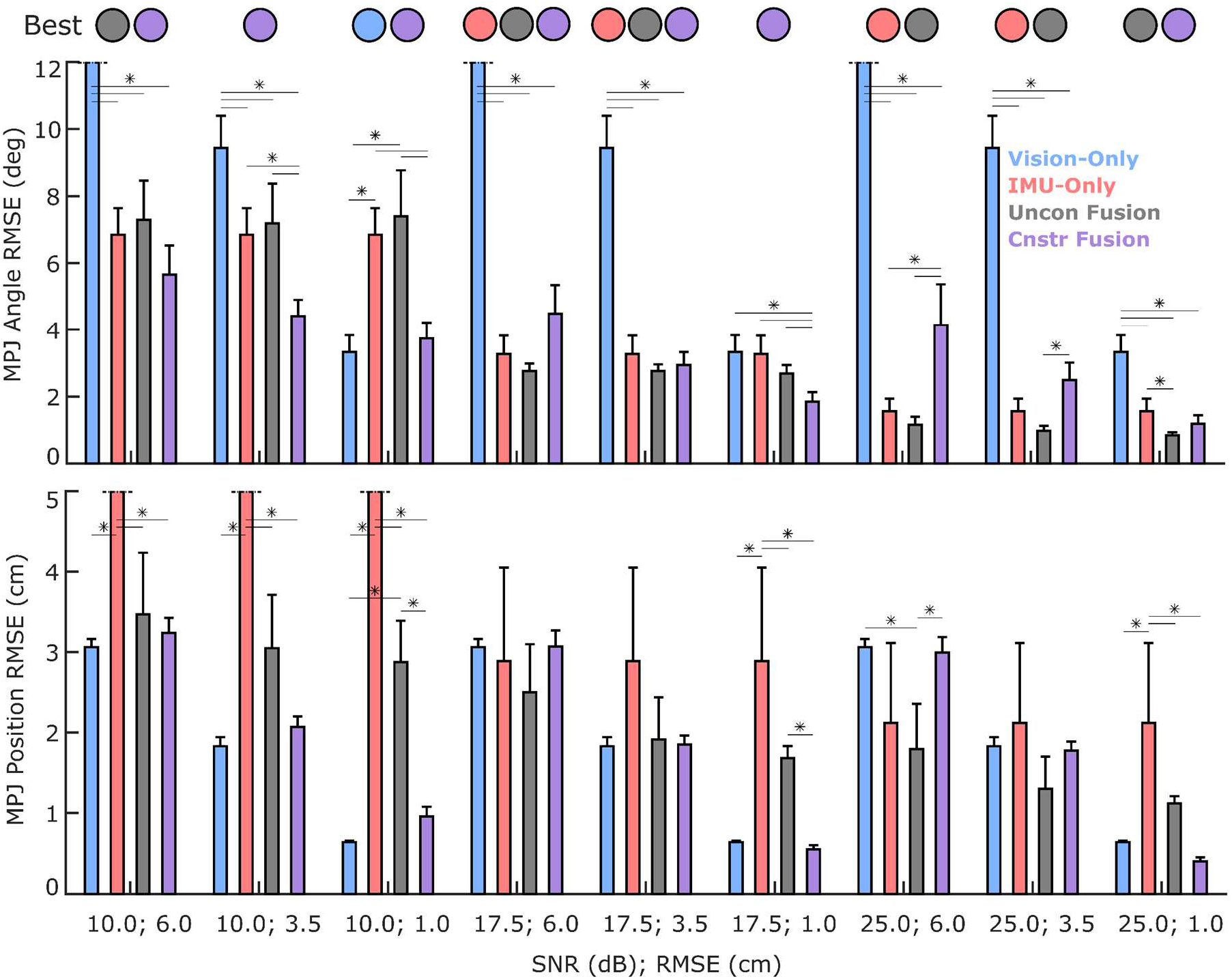
Sensitivity of Markerless Approaches to Noise. Fusion approaches improve results over single modality approaches across almost the entire noise spectrum with few exceptions. Vision-only is consistently outperformed with respect to joint angles, while IMU-only is consistently outperformed with respect to joint center positions. The mean ± standard deviation MPJ angle RMSE (top) and MPJ position RMSE (bottom) show the difference in kinematics predictions across each noise condition for all four techniques (*p < 0.05). The circles indicate which method performed best in each condition, with the color of the circle matching a particular method (blue = vision-only; red = IMU-only; gray = unconstrained fusion; purple = constrained fusion). In the case of multiple circles, two or more methods performed equivalently.

**Fig. 6.**
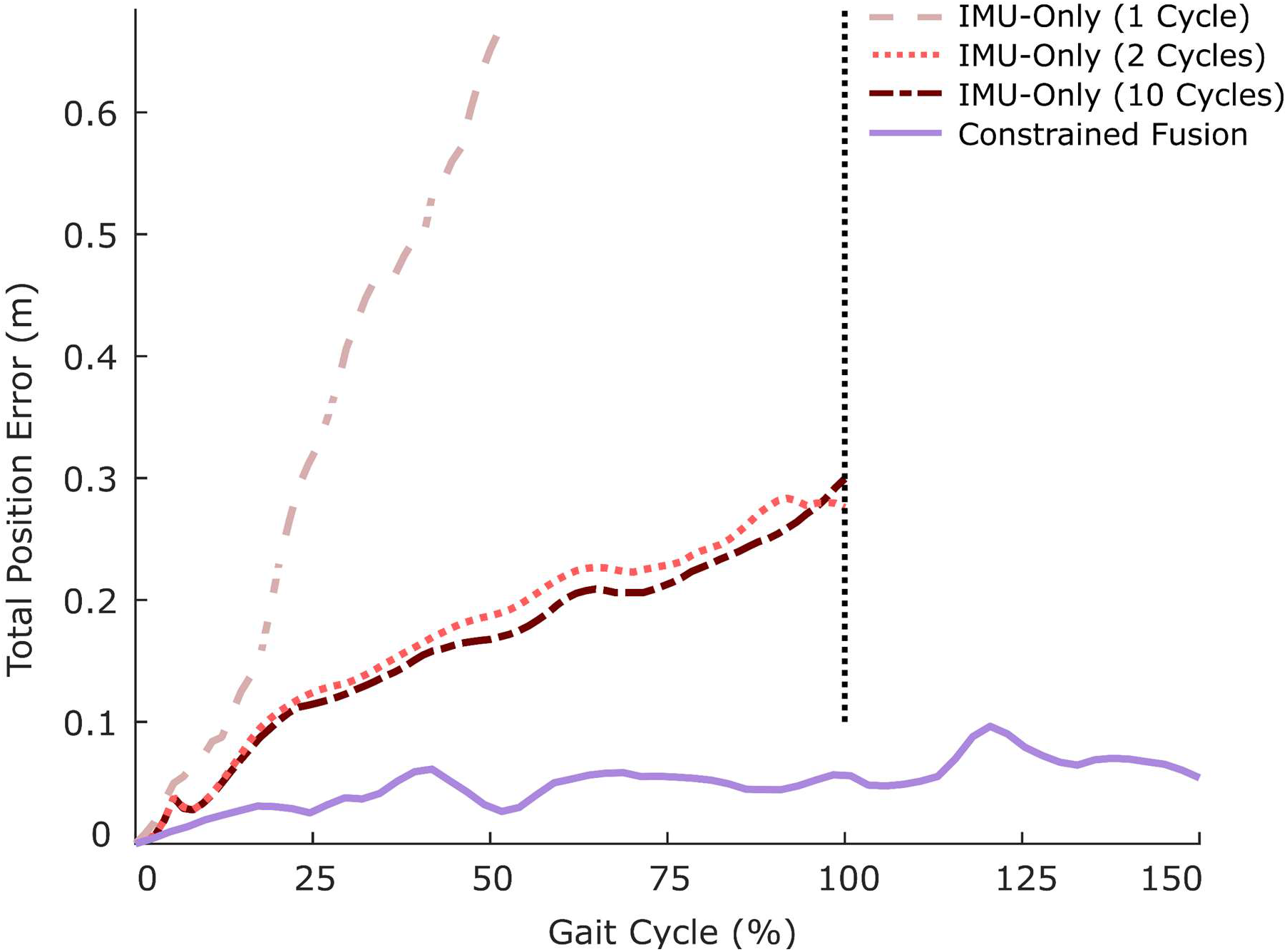
Error Accumulation with IMU Methods. Observing the full body joint center position error over the gait cycle reveals that dynamically constrained fusion and the other techniques eventually reach an equilibrium error, while the IMU-only dynamic optimization continues to accumulate error throughout the simulation duration regardless of the starting IMU data accuracy or the level of denoising. All other approaches can also be run for any arbitrary amount of time, but IMU-only is restricted to complete gait cycles if the periodicity assumption is implemented to reduce drift. However, the rate of error accumulation can be reduced by averaging over multiple periodic gait cycles.

## 4. DISCUSSION

The complementary strengths of wearable sensing, computer vision, and biomechanical modeling could enhance our ability to capture motion and study gait with greater flexibility and cost-effectiveness than current marker-based approaches. Here, we proposed to fuse RGB video and inertial data with a biomechanical model that simultaneously tracks video and IMU data and investigated when this method improves estimation of kinematics over single-modality methods and unconstrained fusion. We found that fusion of video and inertial data improves kinematics over single-modality methods by achieving high accuracy for both joint angles and joint center positions across all the tested video and IMU noise backgrounds. We also found that dynamically constrained fusion with a biomechanical model is advantageous over unconstrained fusion when the quality of inertial sensing data is low and the quality of computer vision models is high, whereas unconstrained fusion is advantageous in the opposite case. When the inertial and vision data noise is equally low or equally high, both types of fusion work equally well, but unconstrained is more computationally efficient.

When interpreting these findings, it is important to consider some of the study’s limitations. Biomechanical modeling simplifications—reducing degrees of freedom, modeling the head, arms, and torso as a single rigid body, and connecting bones to joints by their end points—can affect the results of simulations. Yet, this simplified approach provides baseline insight on how physics-based modeling can contribute to improvement of IMU-video fusion. Our proof-of-concept study provided a strong premise to continue to invest effort in this direction and expand to 3-D models but do so with better awareness of the scenarios under which such computationally expensive approaches are beneficial. We expect that models with greater complexities and constraints, like OpenSim, will amplify but not overturn the conclusions drawn here. Furthermore, we created synthetic data for testing each approach across different noise magnitudes by simply scaling the noise backgrounds inherent to the experimental IMU and video data. We find this approach elegant and the assumption that the noise distribution remains constant across noise magnitudes more reasonable than making assumptions about that distribution (e.g., Gaussian, uniform, etc.), but a validation of the synthetically scaled noise profiles could be used to test that hypothesis in the future. Another limitation is that only walking was considered here. It remains to be determined if the reported findings hold across other activities. As a final note, the video and IMU data may be weighed differently in the cost function based on prior knowledge of their noise profiles, which we did not do. A sensitivity analysis of weights assigned to each modality, however, revealed that this finetuning process does not change the results sufficiently to overturn our primary conclusions (Supplementary Figure 1).

The finding that fusion of RGB video and IMU data is advantageous to single-modality approaches is consistent with findings from other disciplines, despite the lack of exploration in biomechanics. State estimation and simultaneous localization and mapping (SLAM) in autonomous robot navigation is commonly achieved by fusing IMU and video data with extended Kalman filters (R. Smith et al., 1990) and modified particle filters (Montemerlo et al., 2002). Currently, this fusion method provides the most viable alternative to GPS and lidar-based odometry in aerial navigation (Scaramuzza & Zhang, 2020). IMUs overcome visual SLAM limitations like occlusion, motion blur, a lack of visible textures, and inaccurate velocity and acceleration estimates, while videos help enable IMU recalibration in real-time to overcome drift (Mirzaei & Roumeliotis, 2008; Nikolic et al., 2014). The complementary nature of videos and IMUs explains why fusion methods consistently outperformed single-modality methods across the entire range of tested noise conditions and why they should be adopted in biomechanics as they are in robot-state estimation. In biomechanics, fusion can also help overcome soft-tissue motion artifacts since computer vision methods detect joint centers where skin motion is small, whereas IMUs sense inertial changes at the middle of body segments where skin motion is non-negligible. However, while fusion is generally better, attention must be paid to both data quality and computational cost to select the most appropriate fusion approach for a particular application.

The overlap between biomechanical models and IMUs causes the unconstrained and biomechanically constrained fusion approaches to diverge under specific noise conditions. Biomechanical models provide mathematical expressions relating applied forces to rigid-body velocities and accelerations. IMUs provide experimental measurements of rigid-body angular velocities and accelerations. When IMU data are inaccurate, adding a model is beneficial because the underlying optimizer can leverage model physics to reduce dependence on suboptimal IMU data. However, when the IMU data are more accurate than the model due to modeling simplifications, adding the model becomes detrimental. Because IMU data quality is limited by miscalibration errors and soft-tissue artifacts, the incorporation of a biomechanical model will likely remain beneficial for natural environment applications of fusion methods. Furthermore, incorporation of a model is likely to benefit measurements of faster activities associated with larger skin deformations.

As the prevalence of health monitoring in natural environments increases, so will the frequency with which patients and clinicians are charged with setting up lightweight and portable health-monitoring systems. Markerless motion capture methods must therefore be robust to the IMU and camera miscalibrations resulting from suboptimal setups by nonexperts. Since fusion of complementary modalities has proven to be more robust to noisy data than single modality methods, we recommend greater emphasis be placed on thoroughly exploring and benchmarking data fusion approaches for biomechanical applications. Our work provides a preliminary comparison of emerging techniques that could make motion capture more accessible. Our findings could help researchers and clinicians make more informed decisions, weighing the required accuracy for a given application against sensor density and computational complexity. Our published code provides an opportunity to further verify our conclusions with real video and IMU data from different laboratories.

## Acknowledgements

Research reported in this publication was supported by the United States National Science Foundation (NSF) Graduate Research Fellowship Program (award numbers DGE1745016 and DGE2140739), a Korean Government Fellowship, and the NSF Disability and Rehabilitation Engineering program (award number CBET 2145473).

## Data Availability

All the code required to generate the findings of this study is made available via SimTK and GitHub:

https://simtk.org/projects/imuvisbiomech

https://github.com/CMU-MBL/IMUVisionBiomechanics.git

## Conflict of Interest Statement

The authors declare no competing interests.

---

**Supplementary Figure 1.**
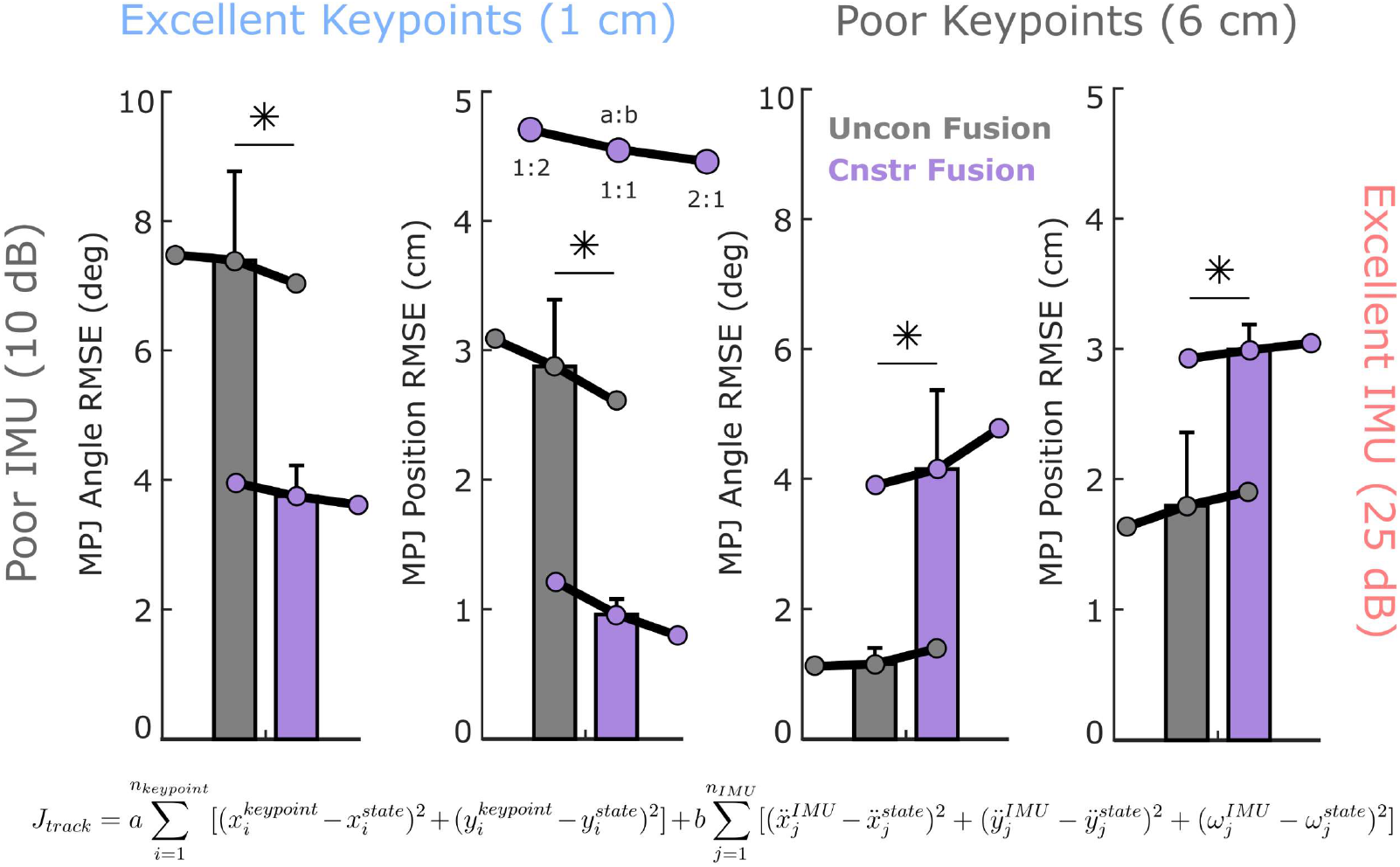
Sensitivity to IMU and Video Data Weights. The cost functions associated with both constrained and unconstrained fusion can use different weights for the video data (a) and IMU data (b). When the noise profile associated with each modality is known, the a:b ratio can be modulated accordingly so that the solution relied more on the higher quality data source. We investigated if fine-tuning of these weights would overturn the conclusions drawn here and found that it would not. Under three scenarios, where a/b = 0.5, a/b = 1, and a/b = 2, the relative performances of the constrained and unconstrained remained the same.

## Notes

https://github.com/CMU-MBL/IMUVisionBiomechanics.git

https://simtk.org/projects/imuvisbiomech

### Competing Interest Statement

The authors have declared no competing interest.

### Summary of Updates

The manuscript was updated to reflect improvements made during peer review with the Journal of Biomechanics

